# Challenges and Solutions in Quantifying Brain β-Hydroxybutyrate (BHB) with ^1^H-MRS Following Oral Keto-Ester Consumption

**DOI:** 10.64898/2026.07.04.736442

**Authors:** Mansimran Virk, Kennedy T. Connors, Razi Kitaneh, Marcella M. Mignosa, Scott McIntyre, Terence W. Nixon, Kelly DeMartini, Stephanie O’Malley, John H. Krystal, Henk M. De Feyter, Gustavo A. Angarita, Graeme F. Mason, Robin A. de Graaf, Chathura Kumaragamage

## Abstract

**Purpose:** β-hydroxybutyrate (BHB), a ketone body and alternative cerebral energy substrate, can be measured in vivo using J-difference edited proton magnetic resonance spectroscopy (^1^H-MRS). Oral ketone supplementation with substrates such as the ketone monoester (R)-3-hydroxybutyl-(R)-3-hydroxybutyrate (KME) and 1,3-butanediol (BD) have gained attention as a mechanism to elevate circulating BHB and induce ketosis without dietary restrictions. Elevated brain ketone availability is of growing therapeutic interest as a strategy to support neuronal energetics in conditions such as epilepsy, neurodegenerative disease, and alcohol use disorder (AUD). However, both pathways introduce BD into the bloodstream, which crosses the blood-brain barrier. Critically, BD exhibits a spectral signature that closely resembles the prominent BHB peak in JDE-MR spectroscopic imaging (MRSI), identified in a pilot AUD study.

**Methods:** Two separate JDE-MRSI acquisitions tailored for BHB and BD editing were implemented, exploiting frequency separation between the BHB (4.14ppm) and BD (3.95ppm) coupling partners of the observed 1.2ppm resonance to independently quantify each metabolite.

**Results:** Brain BD concentrations (0.25-0.58mM) were comparable to or exceeded corresponding BHB concentrations (0.20-0.27mM) in all volunteers after consumption of a single dose of the KME, indicating that BD constitutes a major fraction of the signal conventionally attributed to BHB. Combined BHB+BD concentrations (∼0.45-0.85mM) were consistent with brain BHB values reported in prior studies employing similar doses of the KME, indicating that those measurements likely reflect a combined BHB+BD signal.

**Conclusions:** Separate quantification of the two metabolites is important for interpreting brain ketone studies and for understanding the full pharmacology of KME supplementation.

## Introduction

Under normal physiological conditions, the human brain relies predominantly on glucose to meet the high energy demands of synaptic transmission, ion transport and neurotransmitter synthesis^1,2^. However, when glucose availability is limited (during prolonged fasting, extended exercise, or adherence to a ketogenic diet) ketone bodies cross the blood-brain barrier via monocarboxylate transporters (primarily MCT1 and MCT2) and serve as an alternative energy source^1,3–7^. Among these ketone bodies, β-hydroxybutyrate (BHB) is the most prevalent, capable of supplying up to 60% of the brain’s energy requirement during prolonged fasting^1,3^.

Ketogenic diets, which markedly restrict carbohydrate intake, induce sustained elevations in circulating ketone bodies and have long been used therapeutically in neurological conditions such as epilepsy^8–11^. More recently, increasing evidence suggests that elevated ketone availability may have broader neurobiological benefits in conditions such as neurodegenerative disease^11–15^, cognitive enhancement^11,14,15^, and the treatment of alcohol use disorder (AUD)^16–19^.

Recent studies have shown that exogenous ketone supplementation can rapidly raise circulating BHB levels without dietary restriction or prolonged fasting^1^. Among these, the ketone monoester (R)-3-hydroxybutyl-(R)-3-hydroxybutyrate (KME) is especially effective^20^, producing higher and more sustained blood BHB levels^21^ than ketone salts^22,23^ while maintaining safety and tolerability in healthy adults^5^. These properties make KME a promising candidate for modulating brain energetics in neurological disorders.

Conventional ¹H-MRS quantification of brain BHB is hindered by substantial spectral overlap between BHB resonances and other endogenous metabolites. Robust and non-invasive detection of brain BHB can be achieved using ¹H-MRS, and J-difference editing (JDE)^2,16,24^. In JDE, frequency-selective editing pulses modulate J-coupled spin systems, allowing overlapping signals from uncoupled metabolites to be subtracted while retaining the edited J-coupled resonances of the target metabolite^25^. Several studies have employed this strategy to measure brain BHB following ketone ester ingestion^16,24^. However, accurate quantification can be complicated by a confound that has not been adequately addressed in the literature.

Following oral ingestion, the KME is hydrolyzed in the gastrointestinal tract to yield BHB and BD^4,26^; BD is subsequently metabolized to additional BHB predominantly in the liver and also brain via alcohol dehydrogenase^4,27^ (Figure 1). This two-step pathway produces an immediate and sustained rise in systemic BHB. Critically, BD itself crosses the blood-brain barrier and possesses a spin system that closely resembles that of BHB: its methyl protons resonate near 1.2 ppm are coupled to a methine proton at 3.95 ppm, only ∼0.2 ppm from the BHB coupling partner at 4.14 ppm. When BHB editing targets the 4.14 ppm resonance, the approach can perturb the 3.95 ppm BD coupling partner (based on the editing pulse bandwidth), resulting in a co-edited BD signals at 1.7 ppm and crucially, the 1.2 ppm resonance that overlaps with the target BHB resonance^16,24^. Highly selective editing pulses can reduce this overlap, but magnetic field inhomogeneity will inevitably lead to coedited contributions of BD, which can be especially problematic with MR spectroscopic imaging (MRSI)^28,29^. Reported brain BHB values following KME consumption therefore potentially represent a combined BHB + BD signal, leading to overestimation of true brain BHB concentrations. This is consequential because BHB and BD have distinct physiological effects: ketogenic interventions reduce alcohol withdrawal in humans, an effect attributed to ketone-body action^18,19^, yet BD itself is reported to suppresses ethanol withdrawal at nonintoxicating doses^30^ and produces ethanol-like CNS effects^31^. Without separating BHB from BD, ¹H-MRS cannot distinguish whether observed brain ketone signals reflect BHB-mediated action or BD’s direct pharmacology.

**FIGURE 1.**
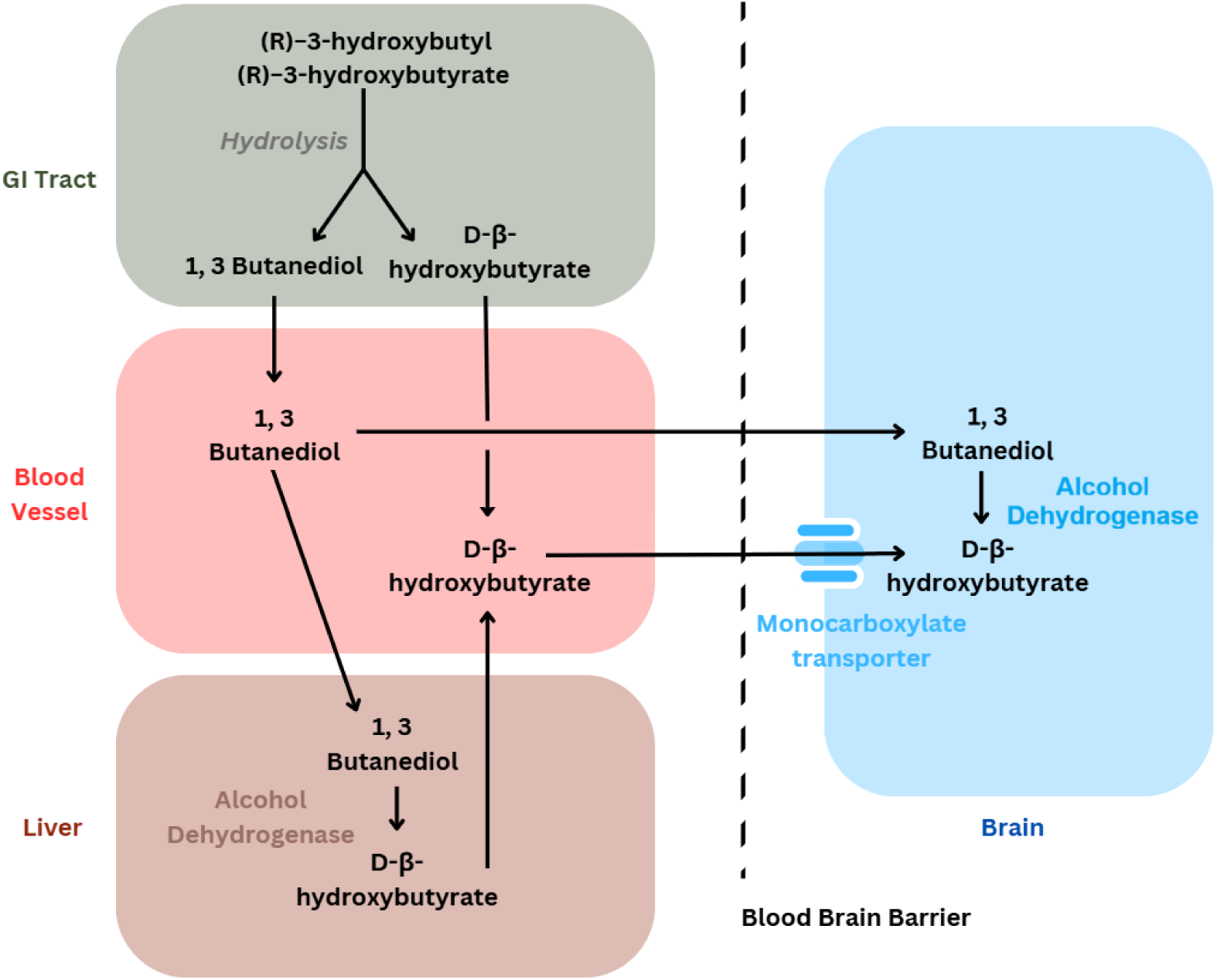
Simplified metabolic pathway of ketone body metabolism following oral consumption of the ketone monoester (R)-3-hydroxybutyl (R)-3-hydroxybutyrate.

To improve the robustness and spatial coverage of BHB detection, this study utilized the MC-ECLIPSE platform^32^, integrating enhanced B₀ shimming via a 54-channel multi-coil array with non-linear gradients to achieve extracranial lipid-suppressed, full-brain coverage within an axial slab for MRSI acquisitions. In a pilot AUD study^33,34^ of brain ketone metabolism, we observed that BD enters the brain at appreciable concentrations following oral KME consumption and contributes substantially to the BHB signal. In follow up BHB-BD differentiation study presented here, we characterize the BD spin system by high-resolution NMR, demonstrate BD co-editing in vivo, develop JDE targeted for BHB and BD for their separate quantification, and evaluate contributions of BHB and BD in the brain. Robust and separate quantification of BHB and BD is achieved by developing distinct BHB-edited and BD-edited MRS experiments that maximize the BHB signal in the BHB-edited scan while strategically minimizing contributions from co-edited BD, and vice versa for the BD-edited scan. We further explore strategies to achieve robust separation of the two metabolites, including simultaneous fitting of BHB-edited and BD-edited experiments.

## Methods

### Study and Participants

This work comprises two studies. The first was an alcohol use disorder (AUD) pilot study (ClinicalTrials.gov Identifier: NCT05937893) investigating brain ketone metabolism following oral KME consumption^33^ (herein called the AUD study). The study included 4 alcohol consumers (AC; 2 males, 2 females; age range: 38-60 years) and 8 healthy controls (HC; 7 males, 1 female; age range: 38-60 years). Full eligibility criteria for both groups are provided in the ClinicalTrials.gov study record^33^. All participants in the AUD study were requested to abstain from alcohol for 48 hours and undergo a 10-hour fast to reduce the impact of ethanol and glucose on keto-ester metabolism. During the AUD study, a confounding co-edited contribution of BD to the edited BHB signal was identified, motivating the second follow up study (herein called the BHB-BD differentiation study). This follow-up study focused on characterizing and separating the BD and BHB signals in 3 separately recruited healthy male volunteers (ages 38-50 years), however since the purpose of the follow up study was to investigate BHB and BD separation, no constraints were imposed on fasting or refraining from alcohol. All procedures were approved by the Yale University Institutional Review Board, and all participants provided written informed consent.

### MR Systems

All MR experiments used a 4 T, 94 cm Medspec scanner (Bruker Corporation, Ettlingen, Germany) equipped with the MC-ECLIPSE^32^ gradient/shim insert and 8-element Tx/Rx volume head coil. High resolution NMR (500 MHz, pulse acquire, 5 kHz BW, 16,384 points, 310 K) of a 1,3-butanediol (BD, Sigma Aldrich, Missouri, USA) sample diluted in phosphate-buffered D_2_O was acquired to characterize the BD spin system (Table 1).

**TABLE 1.**
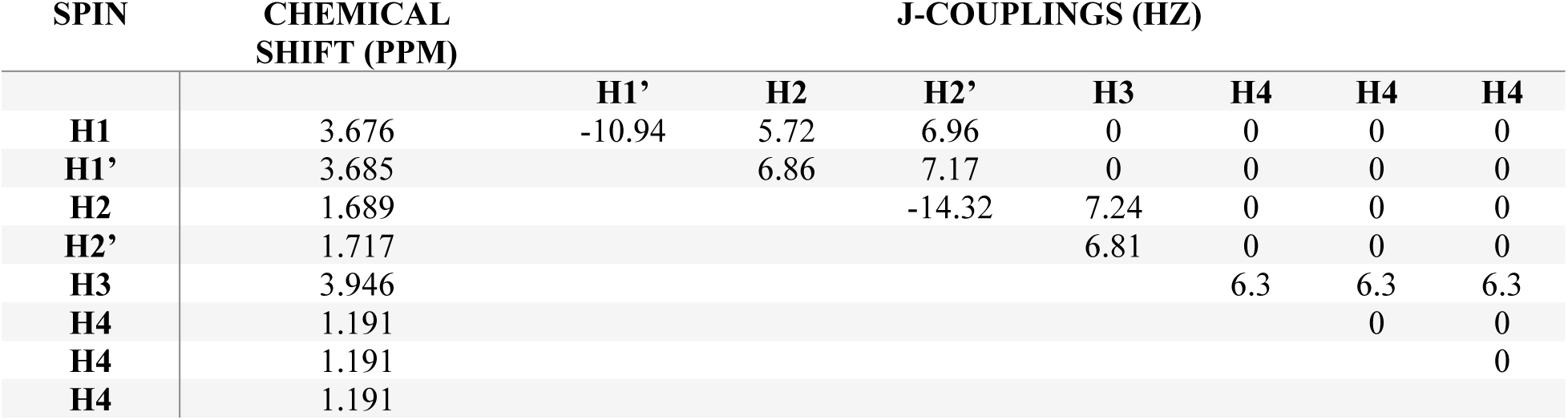
Chemical shifts (ppm) and J-coupling constants (Hz) of the 1,3-Butanediol spin system characterized by high-resolution NMR.

### Study MRSI protocol

A multi-slice gradient-echo MRI (TR/TE = 400/6.5 ms, 180 × 240 × 17 matrix, 1 × 1 × 3 mm^3^ resolution) was used for axial MRSI slab placement, and generating MC-ECLIPSE derived ROIs for extracranial lipid-suppressed MRSI acquisitions^32^. A multi-slice multi-echo B0 mapping method (TR/TE = 90/6.15 ms, 110 × 90 × 17 matrix, 2 × 2 × 3 mm^3^ resolution) was used for B0 shimming over the region of interest.

The MC-ECLIPSE-based MRSI framework^32^ integrating a 4-cycle ECLIPSE outer volume suppression (OVS) module with optimized VAPOR^35^ water suppression was utilized to achieve robust extracranial lipid and water suppression for human brain MRSI. The MC-ECLIPSE-based OVS enables full-axial slice coverage, allowing effective extracranial lipid suppression while preserving cortical regions proximal to the skull. The MRSI acquisition consisted of an asymmetric Shinnar-Le Roux^36^ excitation pulse (Tp = 1.4 ms, BW = 4.2 kHz, asymmetry factor = 0.28), followed by an adiabatic full-passage AFP HS2 (Tp = 2.5 ms, BW = 2.5 kHz) double spin-echo refocusing for axial slab selection. Frequency-selective Gaussian pulses were applied for spectral editing, in principle similar to GABA-edited MRSI previously described^32^. The AUD study utilized Gaussian editing pulses with length of 10 ms (BW = 115 Hz), and the BHB-BD differentiation study Gaussian editing pulses were lengthened to 18 ms (BW = 64 Hz). The JDE-MRSI sequence has an in-plane resolution of 20 x 20 mm^2^ (TR/TE = 2000/144 ms, 9 x 11 matrix, 180 x 220 mm FOV, 2048 complex points, 5.0 kHz sampling BW) over a 20 mm axial slab, leading to an 8 mL nominal volume resolution. The axial slab is placed to encompass prefrontal cortex, anterior cingulate cortex (ACC), and left and right dorsolateral prefrontal cortex (DLPFC) similar to the pilot AUD study. Phase encoding gradients were superimposed on the last spoiler gradient to sample over the elliptical portion of k-space, resulting in an overall duration of 4 minutes 12 seconds for the JDE-MRSI acquisition. A separate water-unsuppressed MRSI scan without editing pulses was acquired (2 minutes 6 seconds) for receiver amplitude/phase correction, and B0 eddy current compensation. An additional low-resolution (4 x 4 x 4 mm^3^) B0 field map was acquired following B0 shimming for post-processing calibrations (detailed below).

The AUD study consisted of a JDE-MRSI acquisition with editing pulses applied at 4.14 ppm and at 5.2 ppm symmetrically around water on alternative acquisitions. The BHB-BD differentiation study consisted of two separate JDE acquisitions to independently quantify BHB and BD, exploiting the small frequency separation between their respective coupling partners. In the BHB acquisition, editing pulses are applied at 4.14 ppm and 3.75 ppm in alternating acquisitions to yield a predominantly BHB-selective edited experiment in which the editing pulses are placed symmetrically around the BD coupling partner to minimize the overall BD contribution, similar in principle to the macromolecular-nulled GABA MRS experiment^37^. The BD JDE experiment employs editing pulses at 3.95 ppm and 4.34 ppm in alternating acquisitions to yield a predominantly BD-selective edited experiment (Figure 2), while minimizing the co-edited BHB contribution due to being symmetrically placed around the 4.14 ppm BD coupling partner. Each JDEMRSI acquisition included two BHB-edited and two BD-edited scans.

**FIGURE 2.**
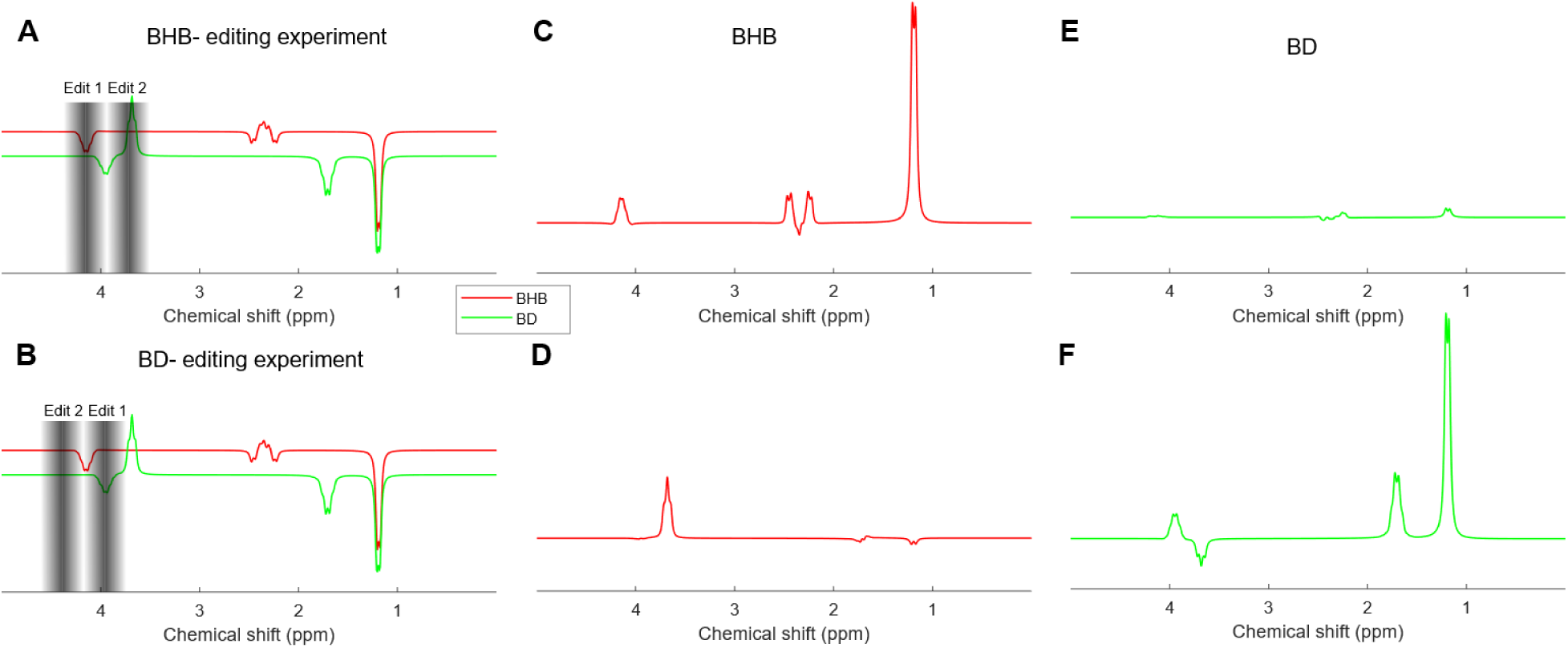
J-difference editing schemes (TE = 144 ms) for BHB and BD. (A) Editing pulses targeted for the BHB-edited experiment: Here Edit 1 is applied on the coupling partner of BHB at 4.14 ppm, and Edit 2 is applied symmetrically around the BD coupling partner at 3.75 ppm to minimize the coedited contribution of BD, while maximizing the BHB contribution. (B) BD-edited experiment: Similar to (A), Edit 1 is applied on the coupling partner of BD at 3.95 ppm, and Edit 2 is applied symmetrically around the BHB coupling partner at 4.34 ppm to maximize BD editing and minimize BHB coediting. The resulting BHB (C-D) and BD (E-F) contributions for the two experiments are illustrated in the second and third columns, respectively considering a 0 Hz B0 offset.

### Magnetic Field Inhomogeneity and Co-editing

Due to the spectral proximity of BD and BHB, each editing experiment in the BHB-BD differentiation study contains signal contributions from both metabolites, with co-edited contributions varying as a function of local B0 field offset (Figure 3). This necessitates a voxel-wise correction using the following system of equations:

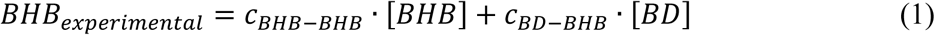

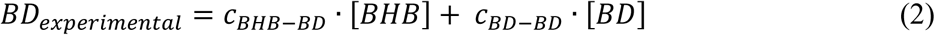

**FIGURE 3.**
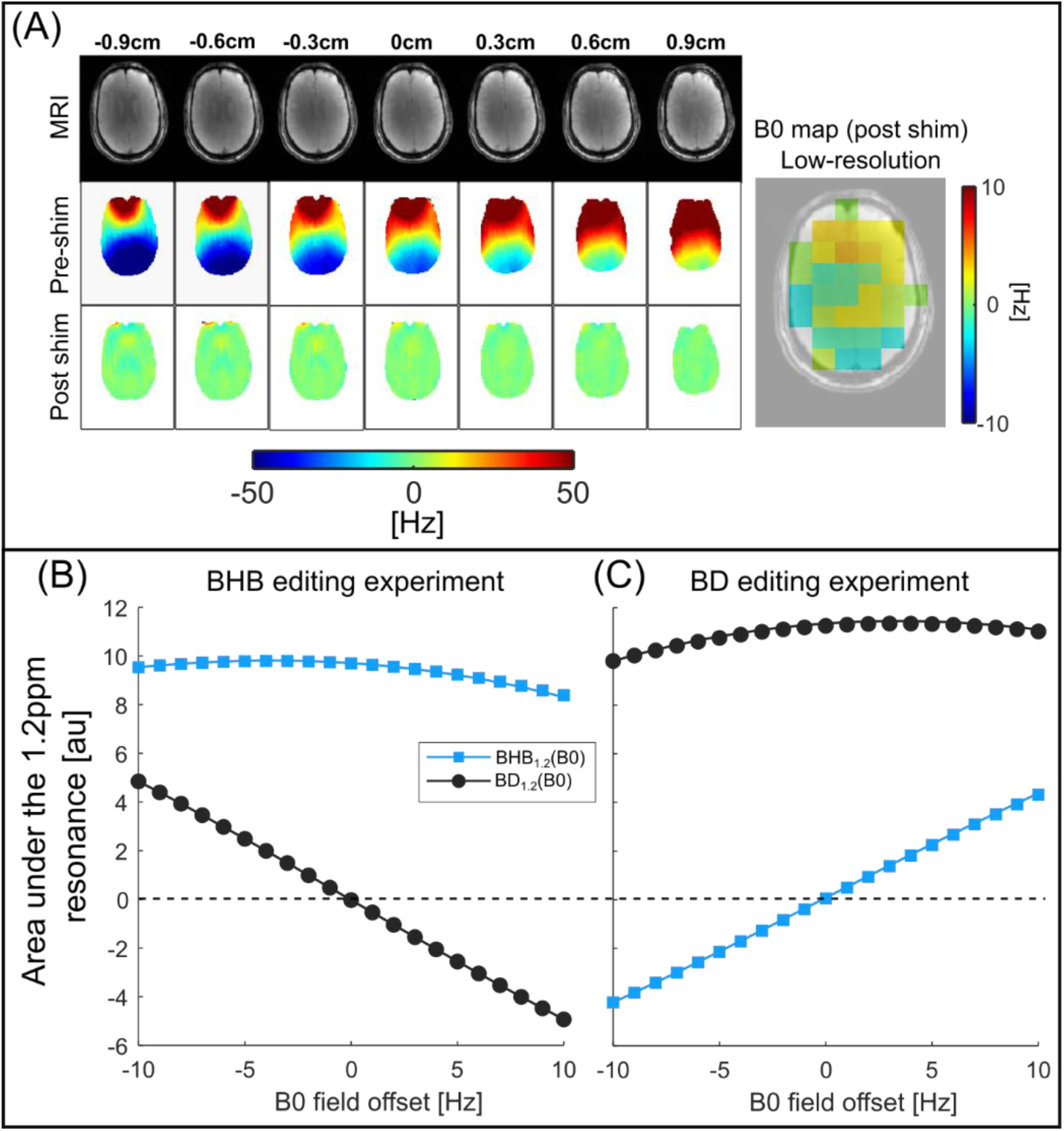
Coedited components of BHB and BD in BHB-edited and BD-edited experiments as a function of B0 field offset. (A) Illustrates a 3x3x3mm3 resolution B0 map covering the 2 cm MRSI slab before and after shimming. On the right, illustrated is a 20x20 mm2 low resolution B0 field map over the MRSI grid constructed from the acquired post-shim B0 map. (B) Simulated contributions of the 1.2 ppm BHB resonance, and co-edited BD resonance as a function of magnetic field offset for the BHB-edited experiment. (C) Simulated contributions of the 1.2 ppm BD resonance, and co-edited BHB resonance as a function of magnetic field offset for the BD-edited experiment.

Here, *BHB_experimental_* and *BD_experimental_* are the concentrations of the 1.2 ppm resonance following LCModel analysis from the BHB-edited and BD-edited acquisitions, respectively. [*BHB*] and [*BD*] are the true metabolite concentrations to be determined. The coefficient *c_BHB_*_−*BHB*_ represents the fractional contribution of BHB in its own experiment given by\ *c_BHB_*_−*BHB*_ = *BHB*_1.2_(*B*_0_)/[*BD*_1.2_(*B*_0_) + *BHB*_1.2_(*B*_0_)] (Figure 3) at the B0 field offset of interest, and *c_BD_*_−*BHB*_ represents the fractional coediting contribution of BD to the BHB experiment *c_BD_*_−*BHB*_ = *BD*_1.2_(*B*_0_)/[*BD*_1.2_(*B*_0_) + *BHB*_1.2_(*B*_0_)]. Similarly, *c_BD_*_−*BD*_ represents the editing efficiency of BD in its own experiment, and the *c_BHB_*_−*BD*_ represents the coediting contribution of BHB to the BD experiment. Coefficients *c* are a function of B0 field offsets and were determined from density matrix simulations considering sequence timings and RF pulse characteristics. The true concentrations [*BHB*] and [*BD*] were determined by voxel-wise inversion of this linear system of equations.

Each JDE-MRSI experiment has a time-resolution of 4-minutes, and averaging of two such scans provided sufficient SNR for metabolite quantification^33,34^. The AUD study consisted of three repeated pairs of JDE-MRSI acquisitions, to enabling time-resolved BHB+ measurements at approximately 55, 75, and 90 minutes following the KME drink consumption (the scan protocol is summarized in Figure S2). In the BHB–BD differentiation study, scan time constraints associated with acquiring both BHB- and BD-edited experiments limited data collection to a single post-supplementation time point for each editing condition. To minimize temporal bias arising from evolving metabolite concentrations, acquisitions were interleaved in the order BHB–BD–BD–BHB. The two BHB-edited and two BD-edited acquisitions were then averaged separately, yielding measurements with effective acquisition times centered at approximately the same midpoint (summarized in Figure S3).

The first MRSI session was acquired prior to ketone supplementation and served to establish baseline BHB+ concentrations in the AUD pilot study, and baseline BHB and BD concentrations in the BHB–BD differentiation study. Next, participants orally consumed a KME drink (deltaG, Orlando, FL, USA) at a dose of 0.4 g/kg body weight (maximum 25 g, corresponding to one serving). Approximately 30-minutes after ingestion, the participant returned to the magnet for subsequent scans. Efforts were made to match the MRSI slab placement for the baseline scan and the second scan. Blood glucose and BHB levels were measured at baseline with a fingerstick sample (Keto-Mojo blood glucose and ketone meter, Napa, CA, USA), and repeat blood BHB measurements prior to and immediately after the second scan session (leading to ∼30-minute and ∼100-minute time points after keto-ester consumption).

### Data Processing and Metabolite Quantification

Spin system simulations, and LCModel (version 6.3-1R)^38^ basis set generation were carried out using SpinWizard^39^, and MRSI data preprocessing was performed in NMRWizard^40^, with both toolkits using MATLAB R2024b^41^. Chemical shifts and J-coupling constants for the BD spin system were estimated using a nonlinear least-squares fit, minimizing the difference between experimental high-resolution NMR data and density matrix-simulated spectra. Spectral fitting and metabolite quantification were performed with LCModel using generated basis sets (including the newly generated BD spin system).

Structural MRI data were reconstructed from raw k-space data and were normalized to unit intensity and exported as NIfTI files. Grey matter (GM), white matter (WM), and cerebrospinal fluid (CSF) segmentations were performed in native space using the Statistical Parametric Mapping toolbox^42^. Default SPM tissue probability maps were employed, with bias regularization (0.001), bias FWHM (60 mm), and native-space probability maps retained for GM, WM, and CSF. Voxel sizes were defined as 1 × 1 mm^2^ in-plane and 6 mm through-plane to reflect acquisition resolution. To account for spatial mismatch between structural and spectroscopic data, the spectroscopic voxel grid was iteratively shifted by ±5 mm in the x- and y-planes until the WM/(GM+WM) weighting on the total Cho concentration was maximized. This x and y offset was applied consistently in all subsequent analyses. For each MRSI voxel, GM, WM, and CSF fractions were estimated by thresholding the corresponding tissue probability maps at 0.5, and a block-averaging approach (20 × 20 pixel blocks) was used to generate coarse voxel-level tissue fractions. GM%, WM%, and CSF% values were exported to CSV format and paired with metabolite quantification outputs.

Metabolite concentrations were normalized to total creatine (tCr), assumed to be 8 mM, and corrected maps were generated for each participant and session. Additional metabolite ratios were calculated, including choline normalized to creatine (Cho/tCr) from unedited spectra, N-acetyl aspartate (NAA/tCr) and glutamate + glutamine (Glx/tCr) from edited spectra. Tissue fractions and metabolite concentrations were pooled across participants to assess tissue-metabolite associations, with regression fits and Pearson correlation coefficients computed for each metabolite as a function of GM and WM fraction.

## Results

### Pilot AUD study

The BD spin system characterized by high resolution NMR is given in Table 1. Figure S1 illustrates simulated spectra at TE = 144 ms (based on BD spin system characteristics in Table 1), demonstrating near-complete overlap of the co-edited ∼1.2 ppm doublet for BHB and BD, along with a secondary BD resonance at ∼1.7 ppm from the H2/H2’ methylene protons. During the AUD pilot study^33^, BHB-edited spectra acquired after KME consumption showed concurrent elevation of 1.2 and 1.7 ppm resonances (Figure S4), consistent with BD co-editing^24^. While the combined

BHB + BD signal (BHB+) can be reliably estimated with high certainty (CRLB values in Figure S4F), separate peak fitting of BHB and BD contributions was unreliable, as evidenced by the fragmented BHB and BD metabolite maps (Figure S4G). This limitation arises from the need to distinguish BD from BHB based solely on the approximately five-fold smaller BD peak at 1.7 ppm (Figure S4D). Figure S5 summarizes the temporal evolution of averaged brain BHB+ concentrations following KME consumption in HC and AC groups derived from Table S2. Relative to baseline, BHB+ levels were elevated at all post-supplementation time points (∼60, ∼70, and ∼90 minutes) in both groups. Time-resolved BHB+ labeling averaged over all voxels was marginally higher for the AC group relative to HC. Statistical testing, however, was not performed owing to the small sample size and the inability to reliably separate BHB and BD contributions to the measured signal.

### Follow up BHB-BD differentiation study

The 64 Hz BW editing pulses used in the BHB-BD differentiation study are sensitive to B0 field inhomogeneity, leading to off-resonance dependent co-editing (Figure 3). By incorporating the B_0_ field offset in each 20 × 20 × 20 mm³ voxel (Figure 3A), the voxel-wise concentrations of BHB and BD were calculated according to Eqs. (1-2), illustrated in Figure 4. Compared with BHB and BD metabolic maps in the AUD pilot study (Figure S4), the combined use of separate BHB- and BD-edited acquisitions with voxel-wise B0-field offset enabled reliable estimation of BHB and BD (Figure 4D–E).

**FIGURE 4.**
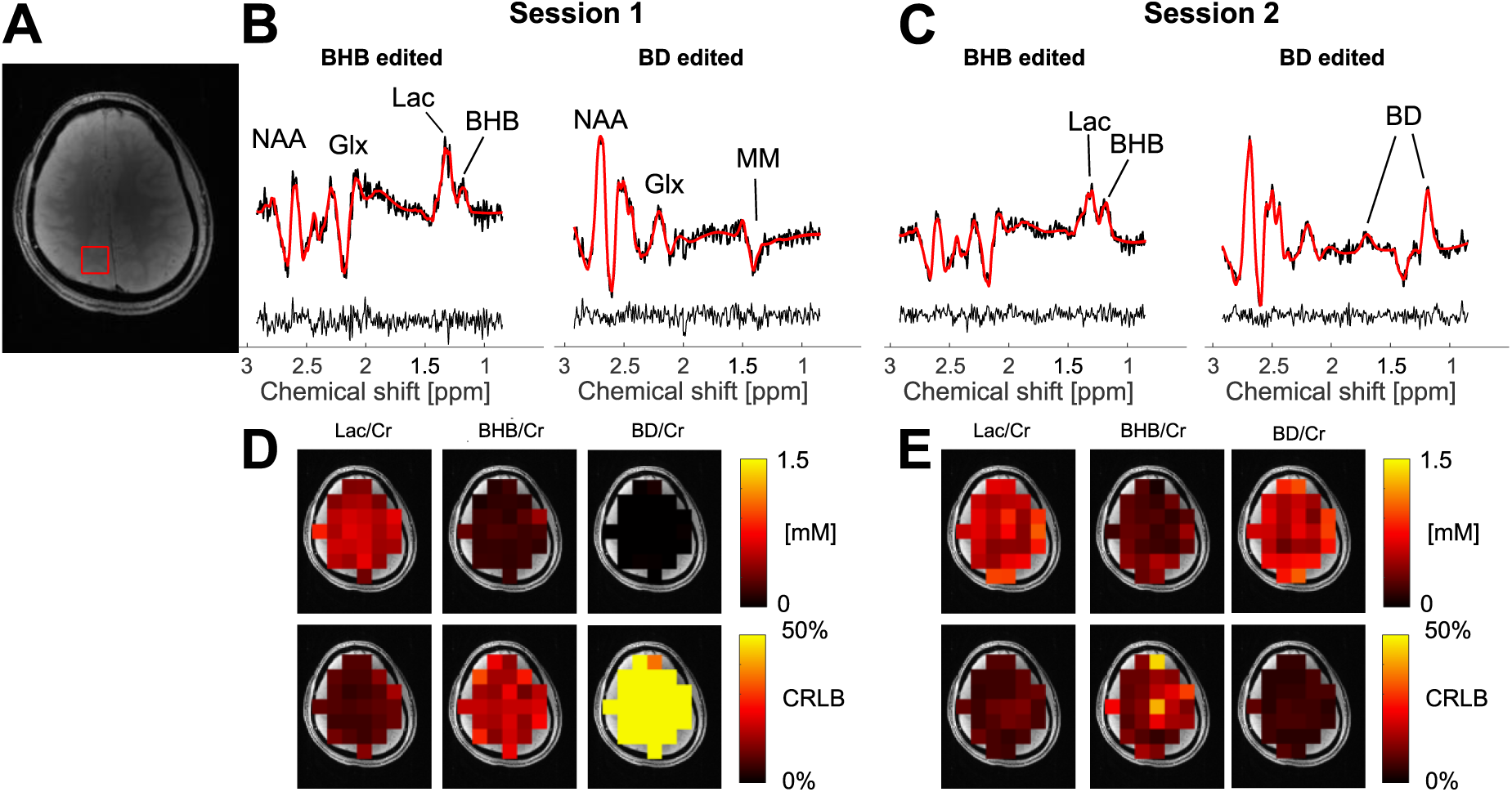
In vivo BHB- and BD-edited MRSI results from a representative participant in the BHB-BD differentiation study. (A) Axial brain image showing voxel placement. (B) BHB-edited and BD-edited spectra for Session 1 (pre-consumption). (C) BHB-edited and BD-edited spectra for Session 2 (post-consumption). (D/E) Metabolic concentration maps (Lac/Cr, BHB/Cr, BD/Cr) and corresponding CRLB maps for both sessions.

Brain BHB and BD concentrations, along with blood BHB measurements for all volunteers, are summarized in Table 2. At baseline, brain BHB concentrations ranged from 0.15-0.18 mM, while BD was negligible (0.0015-0.018 mM) as expected, and blood ketones were 0.1-0.2 mM. Following keto-ester consumption (∼70 minutes post-ingestion), brain BHB concentrations rose to 0.20-0.27 mM, brain BD concentrations rose to 0.25-0.58 mM, and blood BHB levels increased to 2.2-3.6 mM. Notably, post-consumption brain BD concentrations were comparable to or exceeded corresponding BHB concentrations in all three volunteers.

**TABLE 2.** BHB and BD concentrations in the brain (mean ± SD across the axial slice) and blood ketone measurements from all three volunteers in the BHB-BD differentiation study.

|  | BHB_1 | BD_1 | BLOOD<br>KETONES_1 | BHB_2<br>(~70 MIN) | BD_2 (~70<br>MIN) | BLOOD<br>KETONES_2<br>(~90 MIN) |
| --- | --- | --- | --- | --- | --- | --- |
| VOLUNTEER<br>1 | 0.17±0.07<br>mM | 0.006±0.016<br>mM | 0.2 mM* | 0.27±0.1<br>mM | 0.58±0.26<br>mM | 3.4 mM* |
| VOLUNTEER<br>2 | 0.15±0.08<br>mM | 0.0015±0.009<br>mM | 0.1 mM* | 0.2±0.1<br>mM | 0.252±0.16<br>mM | 2.2 mM* |
| VOLUNTEER<br>3 | 0.18±0.1<br>mM | 0.018±0.036<br>mM | 0.2 mM* | 0.21±0.14<br>mM | 0.42±0.22<br>mM | 3.6 mM* |

\*The Keto-Mojo blood ketone meter: ± 0.3 mM accuracy vs laboratory measurements in 90% of samples.

At baseline, BHB showed a weak but significant positive correlation with grey matter fraction (r = 0.25, p = 0.01). Following ketone ester consumption, this BHB-grey matter association was no longer significant (r = 0.08, p = 0.41), while BD showed a significant positive correlation with grey matter fraction (r = 0.19, p = 0.04). As an internal validation of tissue segmentation quality, Cho/tCr showed a strong negative correlation with grey matter fraction at baseline (r = −0.71, p < 0.001), consistent with the expected higher Cho concentration in normal white matter (Figure 5).

**FIGURE 5.**
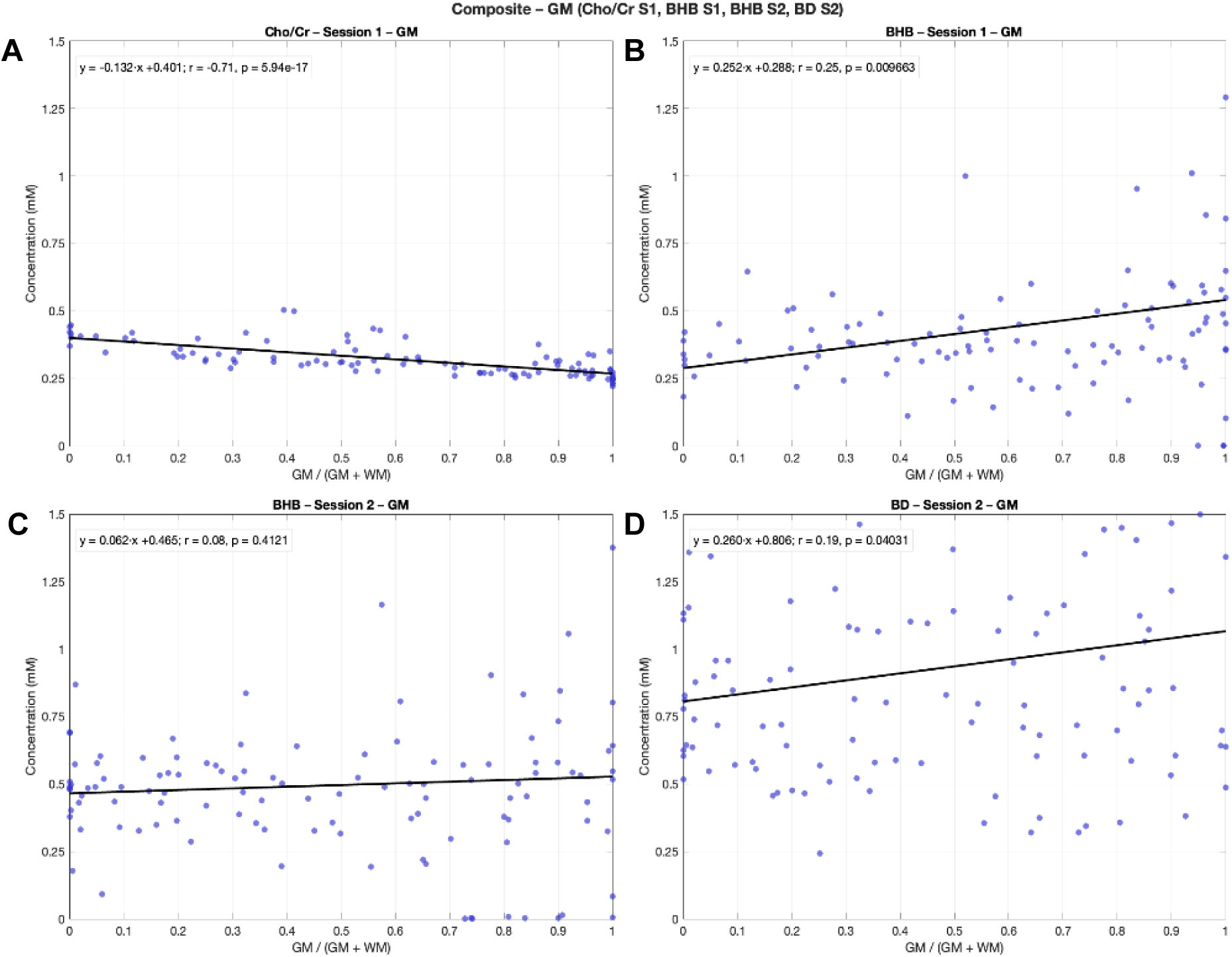
Tissue-metabolite associations. (A) Cho/Cr concentration as a function of gray matter fraction for Session 1. (B) BHB concentration as a function of gray matter fraction for Session 1. (C) BHB concentration as a function of gray matter fraction for Session 2. (D) BD concentration as a function of gray matter fraction for Session 2. Regression lines and Pearson correlation coefficients (r) with P values are shown.

## Discussion

The central finding of this study is that BD accumulates in the brain at concentrations exceeding those of BHB following keto-ester consumption. Furthermore, we demonstrate that BD is co-edited with BHB under standard editing conditions and can contribute significantly to the measured 1.2 ppm signal. Collectively, these findings establish brain BD as a substantial contributor to the post-ingestion KME signal and suggest that BD may contribute to edited ¹H-MRS measurements attributed to brain BHB following ketone ester administration. The ability to distinguish BHB from BD matters because the metabolic fates and physiological roles of BHB and BD differ substantially, with direct relevance for work that models brain energetics or evaluates dose-response relationships using edited ¹H-MRS measurements^3,4^. BD itself is reported to produce ethanol-like CNS depression, induces physical dependence, suppresses ethanol withdrawal symptoms at nonintoxicating doses ^30,31^, and have neuroprotective effects^43^. BD is also metabolized via aldehyde dehydrogenase (ALDH), raising concern for individuals with ALDH deficiency in whom BD clearance may be impaired^44,45^. Consistent with anecdotal alcohol-like sensations reported in humans following BD ingestion^46^, one AUD participant in the AUD study reported a subjective alcohol-like feeling after ketone ester consumption. Although formal neuropsychological testing was not included, these preliminary observations warrant targeted evaluation in future studies.

As shown in the pilot data (Figure S4), the low-SNR 1.7 ppm BD resonance for separation did not achieve physiologically plausible BHB/BD maps. As a potential workaround, all three post-keto-ester JDE datasets (24 minutes of total acquisitions) were averaged and used for spectral fitting. However, the SNR gain was not sufficient to achieve reliable separation. These findings indicate that, within the constraints of typical SNR and scan time with MRSI, conventional JDE may not be sufficient for robust discrimination of BHB and BD in vivo. While improved separation may be achievable with improved SNR, such as with larger voxels in single-voxel spectroscopy, and/or at higher magnetic fields, the proposed JDE approach in the BHB-BD differentiation study provides greater robustness and immunity to B0 variability. However, the ∼16-minute combined acquisition time (two 4-minute scans for BHB editing, and a further two 4-minute scans for BD editing) limits temporal resolution, which may constrain applications requiring repeated measurements for the calculation of dynamic metabolic rates. Future work could explore accelerated acquisition strategies to mitigate this trade-off and investigate optimal scan time windows for BHB signal assessment to support kinetic modelling.

Compared with the BHB–BD differentiation study, the AUD study exhibited higher baseline and post-supplementation BHB+ levels. This difference could be related to differences in monocarboxylic acid transport^47^ or could also result from the standardized 10-hour fast imposed in the AUD study that was not done in the follow up BHB-BD differentiation study, which could increase endogenous ketone production. Baseline differences in blood ketone levels could be masked by the limited accuracy of the Keto-Mojo meter (reported accuracy ±0.3 mM).

The tissue-metabolite associations provide additional support for the BD confound. At baseline, BHB showed a weak positive correlation with grey matter fraction, consistent with the known preferential uptake of ketone bodies in grey matter^3^. Following KME consumption, this BHB-grey matter association was no longer significant, while a BD-grey matter correlation emerged. This pattern is consistent with BD entering the brain in a tissue-dependent manner^3^. The route by which BD crosses the blood-brain barrier is not well characterized; its small molecular size and amphipathic structure are compatible with passive diffusion, but transporter-mediated uptake cannot be excluded^44^.

Beyond these tissue-dependent patterns, much remains unknown about the behavior of BD in the brain following ketone ester ingestion. BD pharmacokinetics in the brain may vary across individuals and with dose; therefore, further studies across a range of KME doses and time points are warranted. More broadly, these findings have practical implications for studies using ¹H-MRS to quantify brain BHB following administration of KME or BD-containing ketone supplements. With conventional single-edited acquisitions, BHB and BD cannot be reliably separated, and the measured signal is most accurately reported as BHB+. In situations in which separate quantification of the two metabolites is needed, the dual BHB-edited and BD-edited acquisition strategy described here provides a route to robust and independent quantification.

In summary, this dual-edit acquisition framework presented allows for independent quantification of brain BHB and BD following oral ketone ester consumption. Given their distinct physiological roles, separate quantification enables study of brain BD pharmacokinetics and supports more accurate interpretation of brain BHB measurements in clinical and research applications.

## ACKNOWLEDGEMENTS

This research was supported by NIH grants R21-EB033911, P50AA012870, R01-EB014861, and NSERC grant RGPIN-2025-05741.

## DATA AVAILABILITY STATEMENT

The AUD study data supporting the findings of this manuscript are currently undergoing curation for deposition and will be made available through the National Institute of Mental Health Data Archive (NDA; https://nda.nih.gov/). Data from the BHB–BD differentiation follow-up study are available from the corresponding author upon request.

## Notes

### Competing Interest Statement

All coauthors except Graeme Mason and John Krystal declare no competing interests.
Graeme Mason reports consulting payments from Leal Therapeutics, Merck & Co., Sumitomo Dainippon Pharma Co., and UCB Pharma for projects unrelated to the present project.
John Krystal has the following conflicts of interest to disclose.
Consultant
Note: The Individual Consultant Agreements listed below are less than $5,000 per year
AbbVie, Inc
Alpha Wave Global
Biogen, Idec, MA
Bionomics, Limited (Australia)
BioXcel Therapeutics
Boehringer Ingelheim International
Cerevel Therapeutics, LLC
Clearmind Medicine, Inc.
Cybin IRL
EpiVario, Inc.
GH Research Ireland, LTD
Janssen Pharmaceutical
Jazz Pharmaceuticals, Inc.
Leal Therapeutics, Inc.
Manifest Technologies
MycoMedica Life Sciences, PBC
Neurocrine Biosciences, Inc
Novartis Pharmaceuticals
Otsuka America Pharmaceutical, Inc.
Praxis Precision Medicines, Inc.
Psylo PTY LTD.
Response Pharmaceuticals, Inc.
Reunion Neuroscience
Spring Care, Inc.
Spruce Biosciences
Takeda Pharmaceuticals, International
Terran Biosciences, Inc.
Tetricus, Inc.
WebMD/ Medscape, LLC
Scientific Advisory Board
Alkermes, Inc
Biogen, Idec, MA
Biohaven Pharmaceuticals
BioXcel Therapeutics, Inc. (Clinical Advisory Board)
Cerevel Therapeutics, LLC
Certigo Therapeutics, Inc
Clearmind Medicine, Inc.
Coalition for Aligning Science
Delix Therapeutics, Inc.
EpiVario, Inc.
Freedom Biosciences, Inc.
Mindset Admin, LLC (DBA Being Health)
MycoMedica Life Sciences, PBC (PMDD Advisory Board)
Neumora Therapeutics, Inc.
Neurocrine Biosciences, Inc.
Novartis Pharmaceuticals Corporation
Praxis Precision Medicines, Inc.
PsychoGenics, Inc.
Response Pharmaceuticals, Inc.
ReST Therapeutics
Spring Care
Takeda Industries
Terran Biosciences, Inc.
Tetricus, Inc.
Stock
Biohaven Pharmaceuticals
Spring Care, Inc.
Stock Options
Biohaven Pharmaceuticals Medical Sciences
Clearmind Medicine, Inc.
Delix Therapeutics, Inc.
Draig Therapeutics
EpiVario, Inc.
Manifest Technologies
Neumora Therapeutics, Inc.
Psylo PTY LTD.
ReST Therapeutics
Spring Care
Tempero Bio, Inc.
Terran Biosciences, Inc.
Tetricus, Inc.
Income Greater than $10,000
Editorial Board, Editor, Biological Psychiatry
Freedom Biosciences, Inc. Science Advisory Board
NON Federal Research Support
AstraZeneca Pharmaceuticals provides the drug, Saracatinib, for research related to NIAAA grant Center for Translational Neuroscience of Alcoholism [CTNA 4]
Novartis provides the drug, Mavoglurant, for research related to NIAAA grant Center for Translational Neuroscience of Alcoholism [CTNA 5]
Cerevel provides the drug PF-06412562 for A Translational and Neurocomputational Evaluation of a D1R Partial Agonist for Schizophrenia (5 U01 MH121766-03)
Patents and Inventions
1. Seibyl JP, Krystal JH, Charney DS. Dopamine and noradrenergic reuptake inhibitors in treatment of schizophrenia. US Patent #:5,447,948.September 5, 1995
2. Vladimir, Coric, Krystal, John H, Sanacora, Gerard Glutamate Modulating Agents in the Treatment of Mental Disorders. US Patent No. 8,778,979 B2 Patent Issue Date: July 15, 2014. US Patent Application No. 15/695,164:
Filing Date: 09/05/2017
3. Charney D, Krystal JH, Manji H, Matthew S, Zarate C., - Intranasal Administration of Ketamine to Treat Depression United States Patent Number: 9592207, Issue date: 3/14/2017. Licensed to Janssen Research & Development
4. Zarate, C, Charney, DS, Manji, HK, Mathew, Sanjay J, Krystal, JH, Yale University Methods for Treating Suicidal Ideation, Patent Application No. 15/379,013 filed on December 14, 2016 by Yale University Office of Cooperative Research
5. Arias A, Petrakis I, Krystal JH. Composition and methods to treat addiction.
Provisional Use Patent Application no.61/973/961. April 2, 2014. Filed by Yale University Office of Cooperative Research.
6. Chekroud, A., Gueorguieva, R., & Krystal, JH. Treatment Selection for Major Depressive Disorder[filing date 3rd June 2016, USPTO docket number Y0087.70116US00]. Provisional patent submission by Yale University
7. Gihyun, Yoon, Petrakis I, Krystal JH Compounds, Compositions and Methods for Treating or Preventing Depression and Other Diseases. U. S. Provisional Patent Application No. 62/444,552, filed on
January10, 2017 by Yale University Office of Cooperative Research OCR 7088 US01
8. Abdallah, C, Krystal, JH, Duman, R, Sanacora, G. Combination Therapy for Treating or Preventing Depression or Other Mood Diseases. U.S. Provisional Patent Application No. 62/719,935 filed on August 20, 2018 by Yale University Office of Cooperative Research OCR 7451 US01
9. John Krystal, Godfrey Pearlson, Stephanie OMalley, Marc Potenza, Fabrizio Gasparini, Baltazar Gomez Mancilla, Vincent Malaterre. Mavoglurant in treating gambling and gaming disorders. U.S. Provisional Patent Application No. 63/125,181filed on December 14, 2020 by Yale University Office of Cooperative Research OCR 8065 US00

