## Supplementary material for "Challenges and Solutions in Quantifying Brain β-Hydroxybutyrate (BHB) with ^1^H-MRS Following Oral Keto-Ester Consumption": Supplimentary material

### Supplementary Figures

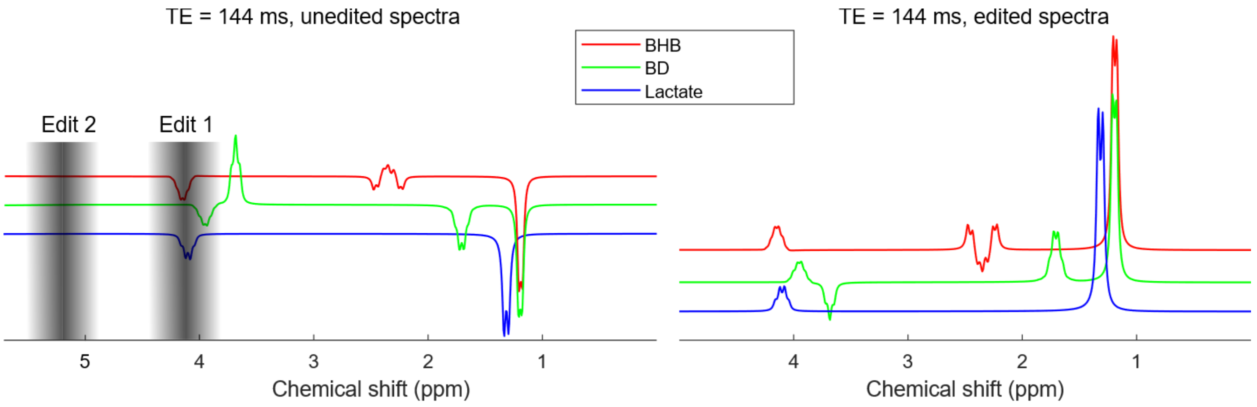


**FIGURE S1**

Spectral footprint of BHB, BD, and lactate without editing (TE = 144 ms, line broadening of 6 Hz) and following J-difference editing (TE = 144 ms, editing pulse 1 applied at 4.14 ppm and editing pulse 2 is applied symmetrically around water at 5.2 ppm). The initial AUD study utilized Gaussian editing pulses (Tp = 10 ms, 115 Hz BW).

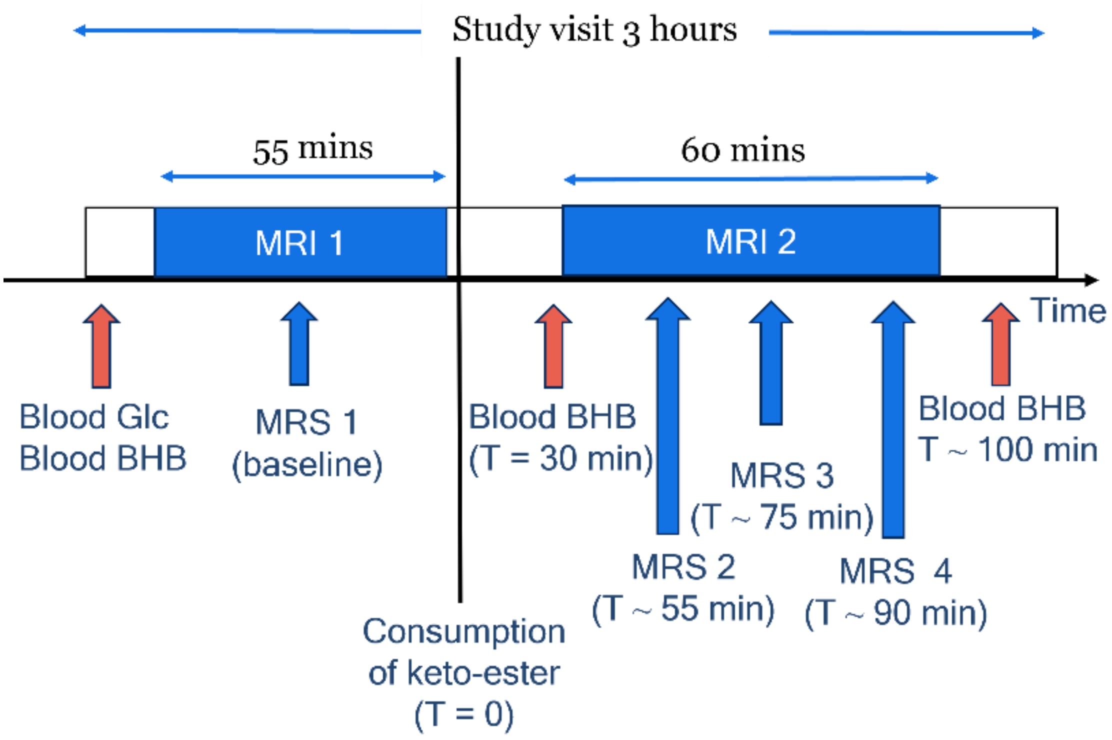


**FIGURE S2**

Study protocol for the pilot AUD study. The MRSI scan protocol consists of two sessions. Baseline BHB edited MRSI is acquired along with anatomical images (~ 55 minutes). Next the volunteer consumes the keto-ester and is placed back in the MRI ~ 30 minutes later for the second session. During this session three 8-minute BHB edited MRSI datasets are acquired to capture BHB+ time points at ~ 55 minutes, ~ 75 minutes, and ~ 90 minutes post-keto ester consumption. The total session two scan time was ~60 minutes in total.

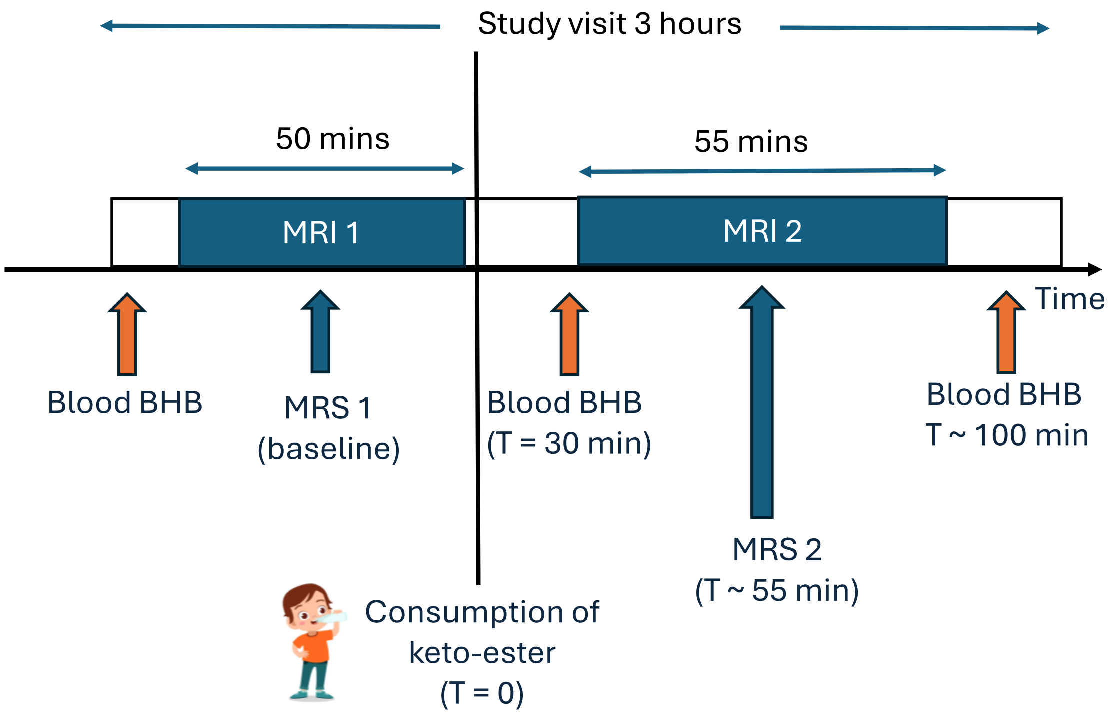


**FIGURE S3**

Experimental protocol and timeline for the follow up BHB-BD differentiation study. The MRSI scan protocol consists of two sessions. Baseline BHB- and BD-edited MRSI scans are acquired along with anatomical images (~55 minutes). The volunteer then consumes the keto-ester and returns to the MRI ~30 minutes later for the second session, during which post-consumption MRSI datasets are acquired (~60-minute mark).

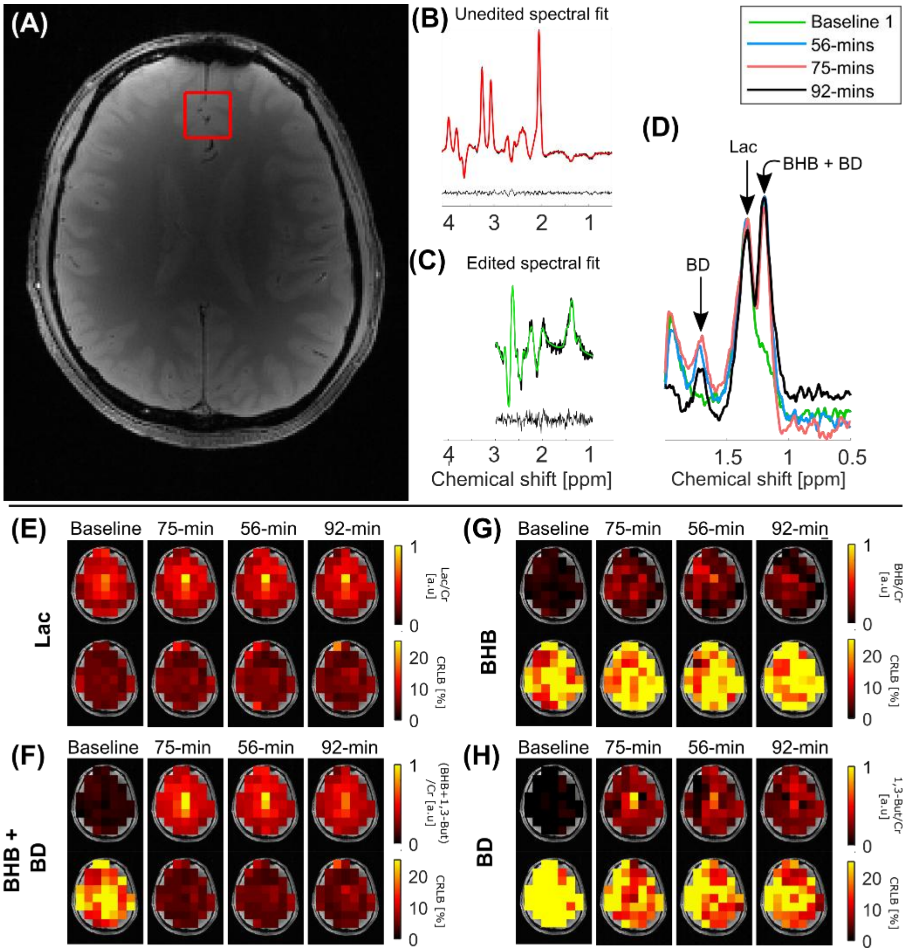


**FIGURE S4**

BHB-edited MRSI acquisition (TE = 144 ms) in a healthy volunteer from the AUD study. (A) Voxel location on anatomical image. (B) Unedited spectrum with LCModel fit. (C) Edited spectrum with LCModel fit. (D) Time course of edited spectra following keto-ester consumption, showing elevated 1.7 and 1.2 ppm resonances corresponding to BD and the combined BHB + BD signal, respectively. Metabolic maps for (E) Lac, (F) BHB + BD, (G) BHB, and (H) BD at four time points (Baseline, 75-min, 56-min, 92-min) with corresponding CRLB values.

**TABLE S1.** Average brain BHB+ concentrations (mean ± SD across the axial MRSI slice) and blood ketone measurements for healthy control (HC) and alcohol consumer (AC) participants in the AUD study. Baseline BHB+ concentrations were obtained during the first MRSI session prior to ketone ester consumption. BHB+ concentrations at time points 1, 2, and 3 (TP1, TP2, and TP3) correspond to the midpoint of the respective MRSI acquisitions during the second (post-consumption) scan session. Blood ketone concentrations were measured prior to ketone ester consumption (baseline), immediately before the second MRI session (~25 min post-consumption), and immediately following the second MRI session (~100 min post-consumption).

| | GROUP | BASELINE BHB+ | TP1 (MIN) | BHB+ @ TP1 | TP2 (MIN) | BHB+ @ TP2 | TP3 (MIN) | BHB+ @ TP3 | BASELINE BLOOD KETONES | BLOOD KETONES (TIME POINT 1) | BLOOD KETONES (TIME POINT 2) |
|---|---|---|---|---|---|---|---|---|---|---|---|
| **1** | HC | 0.37 ± 0.11 | 67 | 1.56 ± 0.48 | 76 | 1.47 ± 0.48 | 97 | 1.39 ± 0.37 | 0.2* | 4.4* (30) | 3.0* (105) |
| **2** | AC | 0.36 ± 0.14 | 60 | 1.62 ± 0.60 | 70 | 1.68 ± 0.60 | 94 | 1.69 ± 0.40 | 0.1* | 4.6* (37) | 2.9* (106) |
| **3** | AC | 0.41 ± 0.13 | 56 | 2.00 ± 0.85 | 66 | 1.88 ± 0.85 | 93 | 1.85 ± 0.60 | 0.1* | 5.0* (25) | 3.7* (102) |
| **4** | HC | 0.46 ± 0.13 | 54 | 2.34 ± 0.76 | 64 | 2.21 ± 0.76 | 84 | 1.96 ± 0.58 | 0.2* | 3.9* (23) | 3.5* (98) |
| **5** | HC | 0.44 ± 0.20 | 54 | 1.83 ± 0.41 | 63 | 1.89 ± 0.41 | 85 | 1.81 ± 0.58 | 0.2* | 4.9* (21) | 2.6* (98) |
| **6** | AC | 0.36 ± 0.13 | 61 | 1.36 ± 0.26 | 70 | 1.70 ± 0.26 | | ABORTED | 0.1* | 1.4* (22) | 4.0* (90) |
| **7** | AC | 0.43 ± 0.14 | 56 | 1.52 ± 0.33 | 65 | 1.55 ± 0.33 | 88 | 1.40 ± 0.57 | 0.2* | 6.6* (20) | 4.0* (98) |
| **8** | HC | FAULT | 52 | 1.28 ± 0.25 | 62 | 1.28 ± 0.25 | 83 | 1.18 ± 0.24 | 0.2* | 4.6* (21) | 4.8* (95) |
| **9** | HC | 0.38 ± 0.14 | 55 | 1.54 ± 0.42 | 64 | 1.51 ± 0.42 | 86 | 1.34 ± 0.47 | 0.1* | 3.2* (23) | 3.1* (97) |
| **10** | HC | 0.39 ± 0.17 | 60 | 1.00 ± 0.40 | 69 | 0.96 ± 0.40 | 91 | 1.24 ± 0.57 | 0.1* | 2.1* (22) | 4.8* (103) |
| **11** | HC | 0.39 ± 0.13 | 58 | 1.55 ± 0.32 | 67 | 1.49 ± 0.32 | | ABORTED | 0.1* | MISSED | 3.0* (93) |
| **12** | HC | 0.41 ± 0.14 | 68 | 1.07 ± 0.35 | 78 | 0.98 ± 0.35 | 96 | 0.91 ± 0.41 | 0.1* | 2.9* (38) | 2.1* (111) |

*The Keto-Mojo blood ketone meter: ±0.3 mM accuracy vs laboratory measurements in 90% of samples.

FAULT – Technical fault of the acquired data.

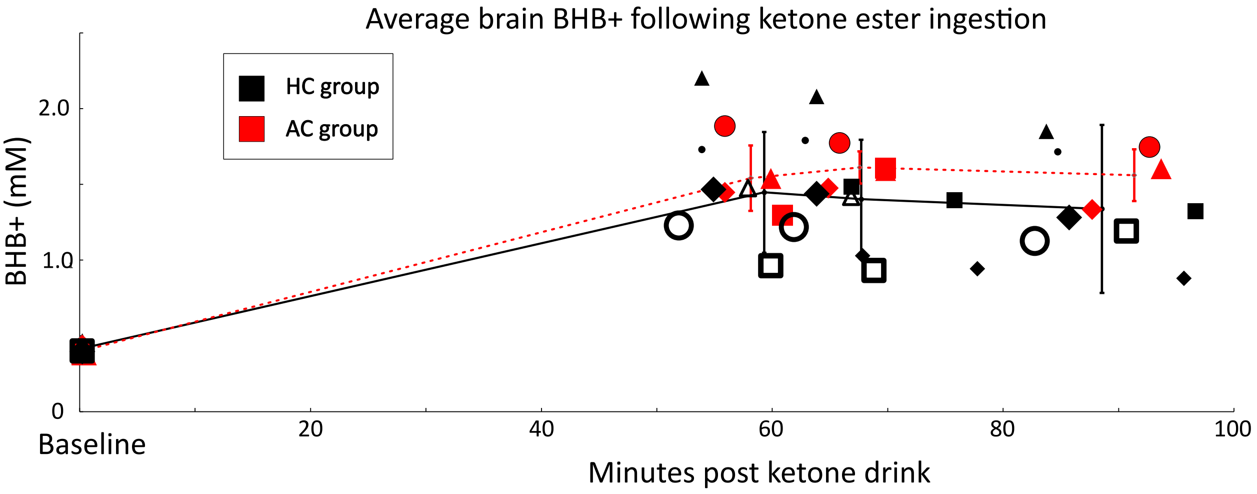


**FIGURE S5**

Averaged BHB+ over the axial MRSI slice for the healthy control (HC) group (n = 8, indicated in black) and alcohol consumers (AC) group (n = 4, indicated in red) from the AUD study. Distinct marker shapes denote individual participants. The first time point at 0 indicates the averaged baseline BHB+ in the brain prior to consumption of the ketone drink. Time point 1, 2, and 3 occurred at ~60 minutes, ~70 minutes, and ~90 minute following oral consumption of the ketone drink during the second session of repeated MRSI scans. The deviation in time for the three time-points across subjects was due to delays in starting the scan session, need for an additional scan etc.  For one HC volunteer, the baseline scan data was unusable due to a technical error, and two volunteer datasets for time point 3 were not acquired (one AC and one HC) due to the need to abort the scan session.
